# Genetic basis of state-dependent courtship sounds in inbred mice

**DOI:** 10.1101/2024.07.22.604528

**Authors:** SaeYeon Na, Jia Ryoo, Chang Bum Ko, Seahyung Park, Daesoo Kim

## Abstract

Courtship behaviors consist of two phases, namely the appetitive and consummatory states. Despite the long history of the concept, few studies have been done regarding the genetic contribution on those two different phases. Male mice are known to produce distinct ultrasonic vocalizations (USVs) as they progress through courtship states^1-14^, and these courtship sounds also show strain-specific differences^15-18^. Here, we delved into USV syllable patterns emitted during specific courtship actions using inbred mouse strains and their progeny: C57BL/6J (B6) mice, 129S4/SvJae (129) mice, and their second filial generation (F2) offspring of mixed genetic backgrounds. B6, 129, and F2 mice generated similar USV syllables during mounting behavior. In contrast, B6 and 129 mice showed different USV syllable patterns during body and anogenital sniffing behavior, and the USV syllable usage of F2 mice in this courtship state diverged according to the degree of genetic similarity with B6 or 129 mice. From these results, we propose that differential selection pressures^19-20^ favored diversity in appetitive behavior but conservation in consummatory behaviors.

## RESULTS AND DISCUSSION

### USV syllable usage can exhibit genetics-related diversity or homogeneity during courtship actions

The courtship behavior of male mice starts from the appetitive state, during which body and anogenital investigation is used to recognize and select an appropriate mate. This is followed by the consummatory state, which starts with a series of mount attempts^1-14^. Since male mice are known to produce specific ultrasonic vocalizations (USVs) during appetitive and consummatory actions, we aimed to identify the effect of genetics on these state-dependent USVs. We used the VocalMat software^21^ to identify the action-dependent USV syllables of C57BL/6J (B6) mice, 129S4/SvJae (129) mice, and their F2 progeny in the presence of freely moving female intruders (Figure 1). We isolated 12 classical USV syllable types as previously described^15,18,22^ and excluded Noise from further analysis (Figure 1A).

**Figure 1.**
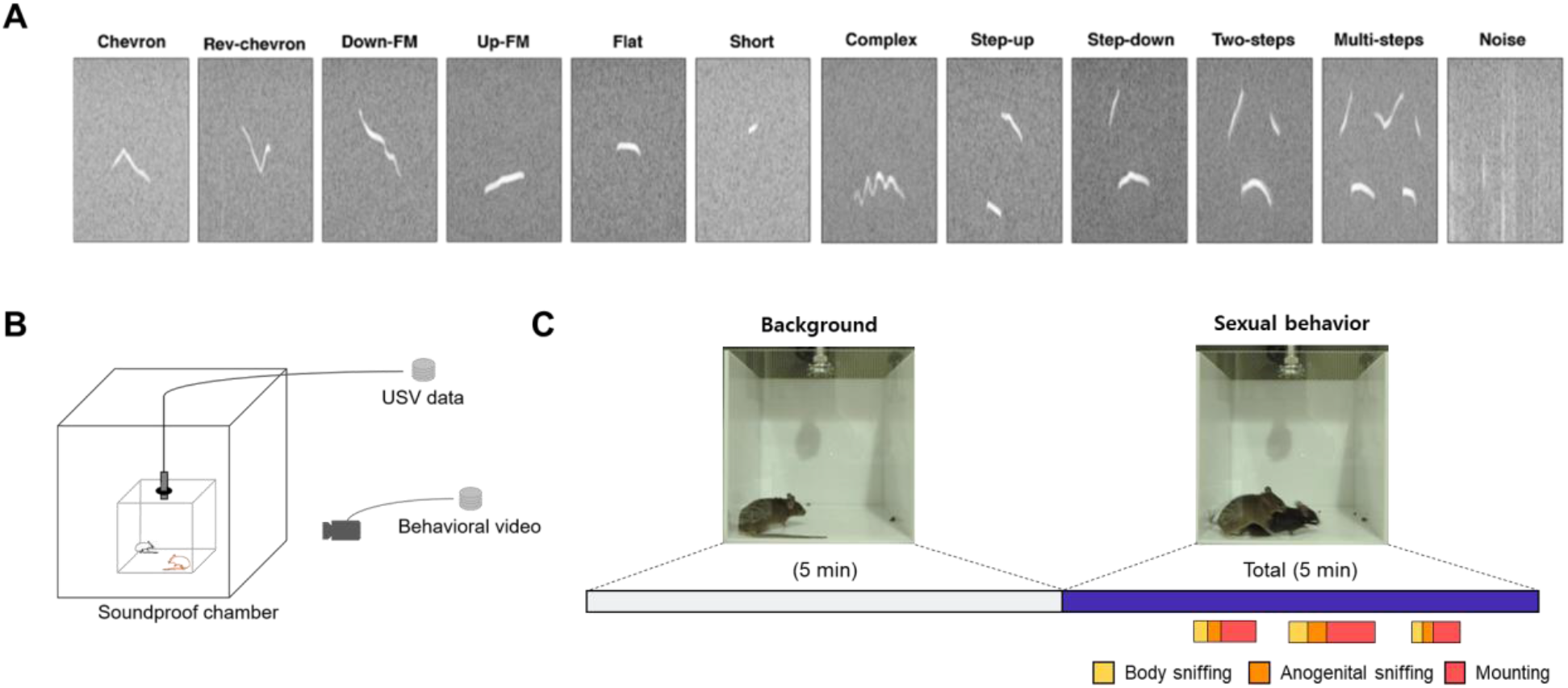
Experimental scheme. (**A**) Syllable classification of Chevron, Reverse chevron, Down-frequency modulation, Up-frequency modulation, Flat, Short, Complex, Step-up, Step-down, Two-steps, Multi-steps, and Noise from VocalMat results. Images were taken from Fonseca et al.^21^ (**B**) Experimental design. (**C**) Representative sample video frames from background and sexual behavior recording. The colors indicate the actions for which the POs of each syllable type were calculated and were as follows: violet, total duration; yellow, body sniffing; orange, anogenital sniffing; red, mounting.

Since the genotype of F2 mice depends on contributions from both 129 and B6 strains, direct comparison of USV syllable usage between F2 and parental strains is statistically inappropriate^23^. Instead, we devised a genetic similarity score that reflected the relative genetic proximity of a given F2 individual to the 129 or B6 strains. The genetic similarity score was calculated using Chevron and Up-frequency modulation (Up-FM) syllables, which are known to be dominant USV syllable markers for B6 and 129 mice, respectively^17,18^. The genetic similarity score was designed to be higher when the USV syllable-related genetics were closer to 129 and lower when they were closer to B6.

To examine how syllable usage varied among animals of different genetic backgrounds, we examined the probabilities of occurrence (POs) for nine syllable types (from among the 12 syllables, Chevron, Up-FM, and Noise were excluded) during distinct actions of sexual behavior (Figures 1B and 1C).

Comparison of syllable usage between B6 and 129 mice during the total duration revealed significant differences (p<0.05) across various syllable x strain pairs (p<0.00002, two-way mixed ANOVA, Table S1), underscoring the presence of strain-specific vocalization patterns.

To test whether the syllable usage over the total duration of courtship behaviors varied proportionally by the genetic similarity with a constant rate of change, we included F2 mice and assessed whether there was a linear relationship between genetic similarity and syllable usage. We employed a general linear model with the genetic similarity score of each mouse and the first principal component (PC1) from the principal component analysis (PCA) results obtained from the nine assessed syllables (Figure 2A). This regression identified a statistically significant linear relationship (p<0.05) between the genetic similarity score and the dimensionality-reduced overall syllable usage (p=0.0005, Figure 2B, Table S3). In other words, the B6, 129, and F2 genetic similarity scores and the summarized syllable usage exhibited a linear relationship during the total sexual behavior period.

**Figure 2.**
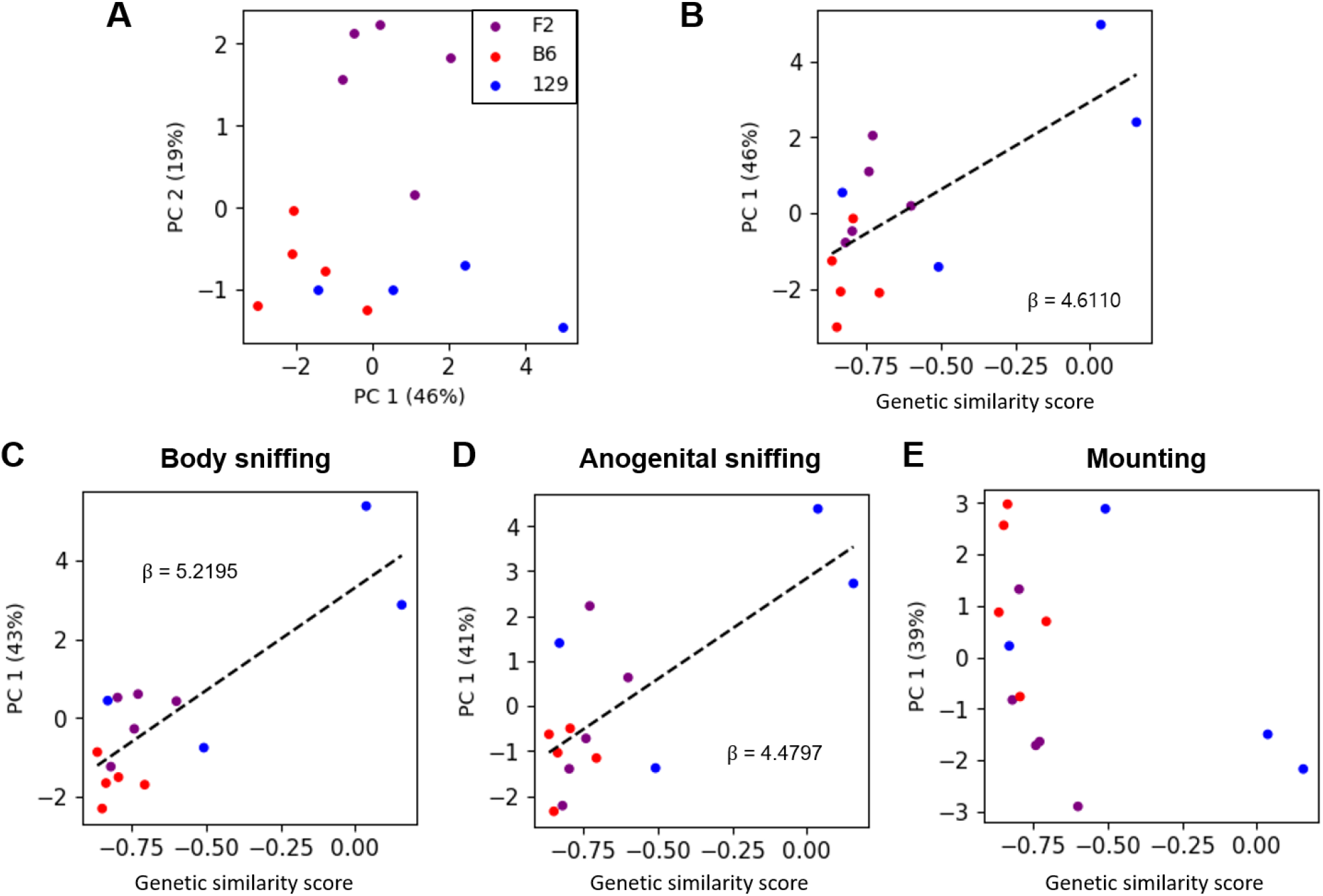
The genetic similarity score exhibits linear relationships with the dimensionality-reduced syllable POs for total duration, body sniffing, and anogenital sniffing. (**A**) Scatter plot of the first and second principal components of the POs for syllables. PC1 explained about 46% of the variance and PC2 explained about 19%. Scatter plot of the genetic similarity score and PC1 values for (**B**) total duration, (**C**) body sniffing, (**D**) anogenital sniffing, and (**E**) mounting. Each point represents an individual animal, with genotypes indicated by the color as follows: purple, F2 (n=5); red, B6 (n=5); and blue (n=4). A Gaussian family generalized linear model with identity link function was employed; Bonferroni correction was utilized for the analyses presented in C, D, and E. Dashed line indicates p<0.05.

To assess whether the strain-specific syllable usage was associated with specific mating-related actions, POs were calculated in three actions to capture the difference in appetitive (body sniffing and anogenital sniffing) versus consummatory (mounting) action. To first assess for syllable usage difference between the B6 and 129 strains, we compared the POs of these strains for each action. We found that B6 mice and 129 mice showed significant differences (p<0.05, Bonferroni correction) in the interaction of strain and syllable during the body sniffing action and the anogenital sniffing action, but not the mounting action (two-way mixed ANOVA, body sniffing: p<0.00005; anogenital sniffing: p<0.000006; mounting: p>0.4; Table S1). The regression results showed that there was a significant linear relationship (p<0.05, Bonferroni correction) between the genetic similarity score and PC1 during the body sniffing action and the anogenital sniffing action, but not the mounting action (body sniffing: p<0.0000006, anogenital sniffing: p<0.0006, mounting: p>0.3, Figure 2C-2E, Table S3). Thus, we observed a between-strain difference and linear relationship in the appetitive stage of sexual behavior (body and anogenital sniffing action), but not the consummatory stage (mounting).

### Two-steps and Flat syllable usage diversity during sniffing

To elucidate which syllable(s) largely contributed to the pattern of strain-specific syllable usage in the actions prior to mounting, we conducted post-hoc tests. Tukey’s HSD (honestly significant difference) test was applied between strains for the PO of each syllable in each action. The POs of Down-FM and Short were higher in B6 mice during the total, body sniffing, and anogenital sniffing actions; that of Two-steps was higher in 129 mice during the total, body sniffing, and anogenital sniffing actions; and that of Flat was higher in B6 mice during the total and body sniffing actions. The POs of the remaining syllables did not show any significant difference between B6 and 129 mice (Tukey’s HSD test, Figure 3A-3C, Table S2). During the appetitive state of sexual behavior, B6 mice used more Down-FM and Short syllables, while 129 mice used more of the Two-steps syllable. During body sniffing, B6 mice dominantly used the Flat syllable.

**Figure 3.**
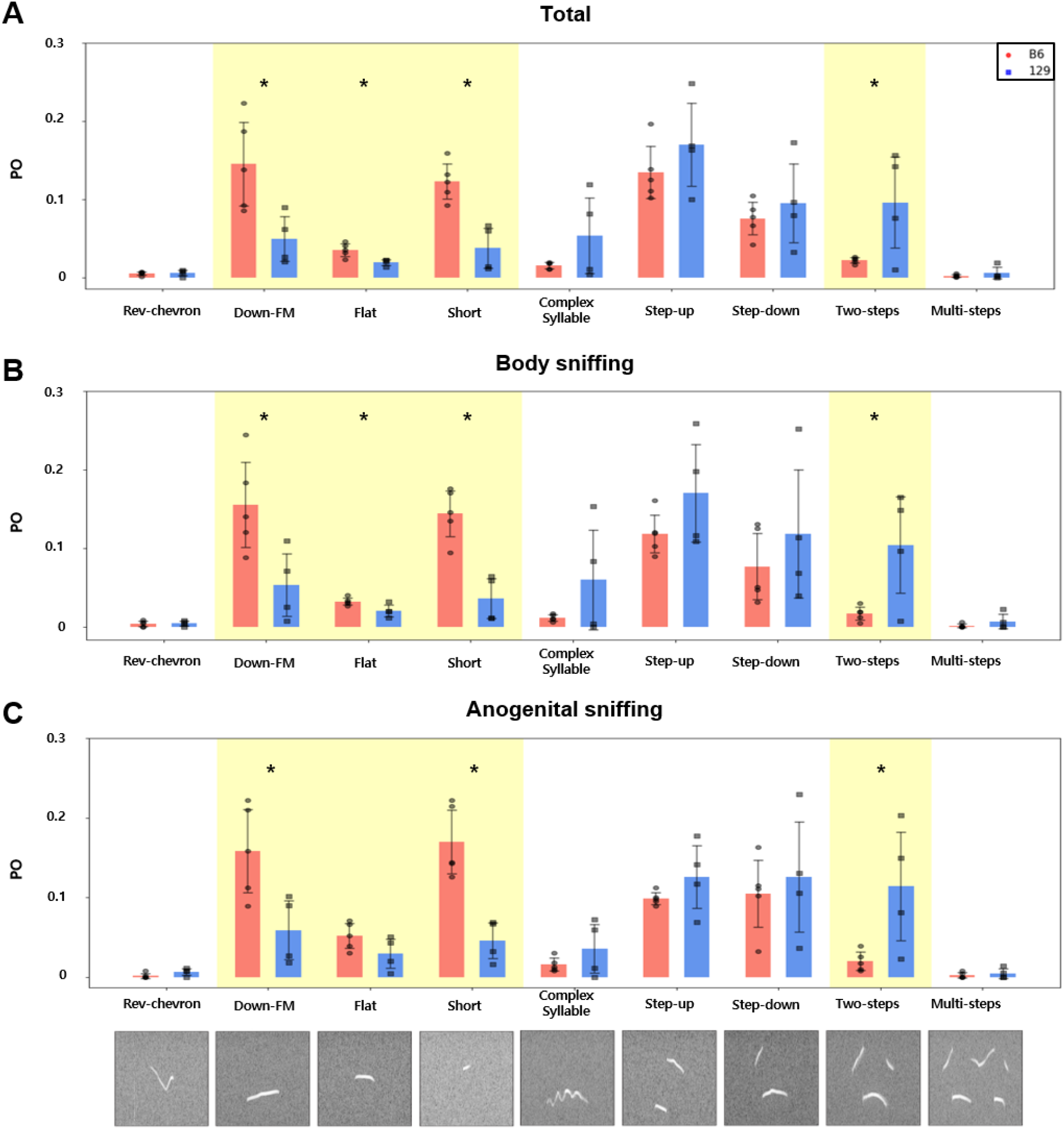
POs differed significantly between B6 and 129 strains for Down-FM, Flat, Short, and Two-steps for the total duration and the different action periods. The between-strain differences in POs were compared during (**A**) the total duration and the (**B**) body sniffing and (**C**) anogenital sniffing actions. Each black point represents the PO of that syllable in each male mouse (B6, circle, n=5; 129, square, n=4). Each red and blue bar represents the mean POs of B6 and 129 mice for each syllable. The yellow background represents the syllables with POs that differed significantly between B6 and 129 mice in at least one action. All data are presented as mean ± SD. *p < 0.05.

The general linear model was fitted to the data with the genetic similarity score used as the independent variable and the PO of each syllable used as the dependent variable, with the goal of identifying syllables that showed linear relationships when we compared the similarity score with the total duration and the body sniffing and anogenital sniffing phases (Figure 4). Bonferroni correction was applied to determine the significance (p<0.05) of linearity. The POs of the Complex and Two-steps syllables showed positive linear relationships with the genetic similarity score in all three actions; that of the Short syllable showed a negative linear relationship with the genetic similarity score during the total duration but not the other actions; and those of the Step-up (positive) and Flat (negative) syllables exhibited positive and negative linear relationships, respectively, during the body sniffing action (Figure 4, Table S3).

**Figure 4.**
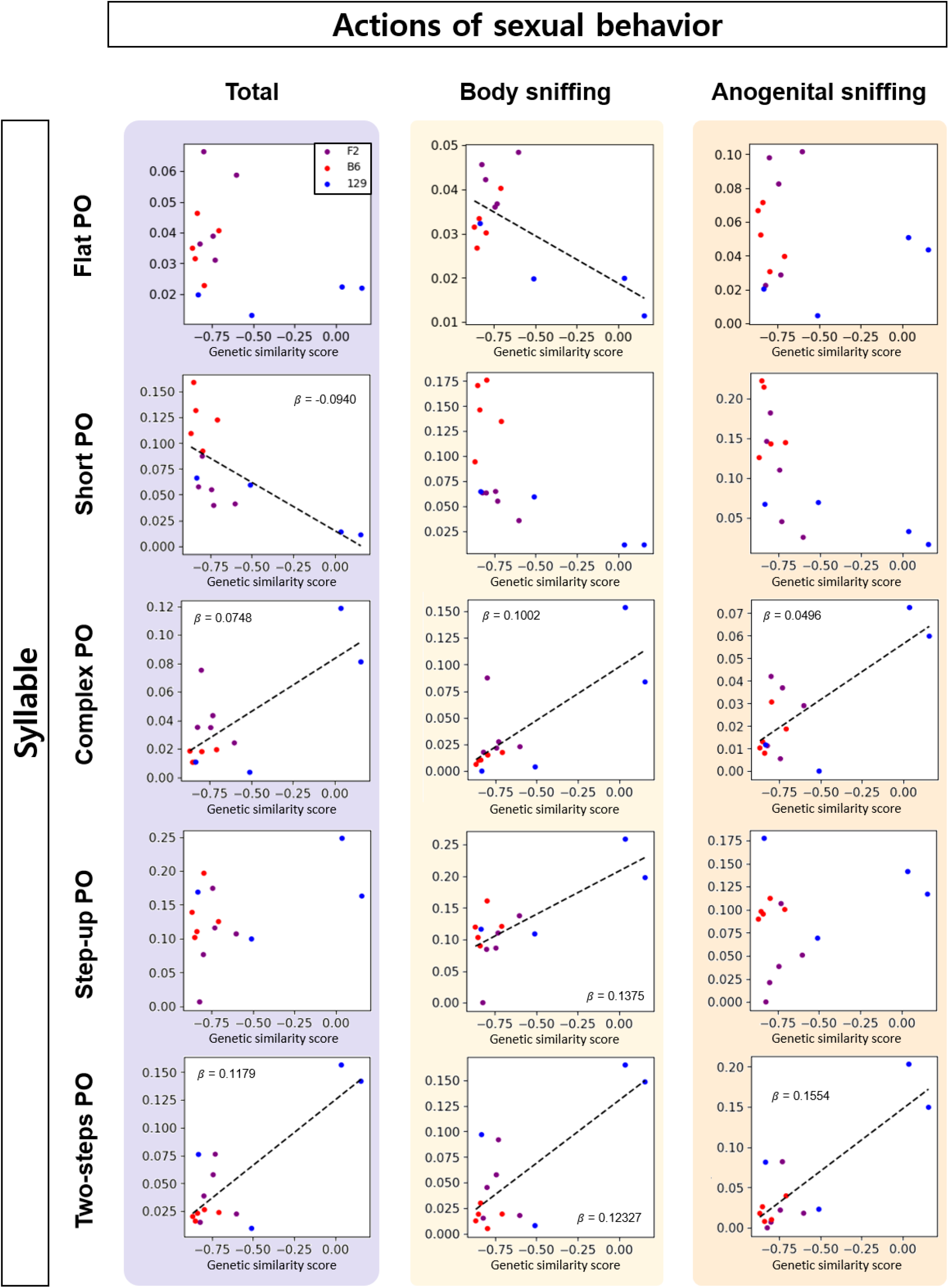
The POs of the Flat, Short, Complex, Step-up, and Two-steps syllables are proportional to the genetic similarity score with a constant rate of change. During the total duration (light violet), the POs of the Short, Complex, and Two-steps syllables showed significant linear relationships with the genetic similarity score; during body sniffing (light yellow), those of the Flat, Complex, Step-up, and Two-steps syllables showed this relationship; and during anogenital sniffing (light orange), those of the Short and Two-steps syllables showed this relationship. Each point represents an animal; purple, F2 (n=5); red, B6 (n=5), and blue, 129 (n=4). A Gaussian family generalized linear model with identity link function was applied, and Bonferroni correction was used. Dashed line indicates p<0.05.

Next, we assessed the linearity between the usage of specific syllables and the genetic similarity score of B6, 129, and F2 mice. The Complex and Two-steps syllables exhibited positive linear relationships with the genetic similarity score, suggesting that they represented 129-like syllables. In other words, male F2 mice that were genetically closer to the 129 strain used more Complex and Two-steps syllables during mating, particularly during the body and anogenital sniffing actions, whereas those closer to the B6 strain used fewer of these syllables during these actions. The Step-up syllable showed a similar pattern, but only during the body sniffing action. The Short and Flat syllables exhibited negative linear relationships with the genetic similarity score, suggesting that they represented B6-like syllables. In other words, male F2 mice that were genetically closer to the B6 strain used more of the Short syllable during the total duration and more of the Flat syllable during the body sniffing action, while those closer to the B6 strain used fewer of these syllables during these actions.

### Genetic basis for USV syllable usage diverging during the appetitive stage and remaining homogeneous during the consummatory stage

Our study delved into how genetic background influences ultrasonic vocalization (USV) syllable usage during various courtship actions in inbred male mice. Studies have shown that genetic background and behavioral state^15^ are the primary factors of syllable syntax. However, the interplay between these factors has not yet been studied in depth. We found that, in sexual behavior of male mice, USV syllable usage differed by the genetic background during appetitive actions but not consummatory actions. B6 mice, 129 mice, and their F2 descendants produced similar syllable patterns during mounting but not during appetitive actions.

Specifically, the Flat and Two-steps syllables demonstrated strain-dependent usage during the appetitive phase. B6-like F2 male mice used more of the Flat syllable and 129-like male mice used more of the Two-steps syllable. These syllables were distinct from those exhibiting strain-specificity throughout the entire course of sexual behavior, as well as from those primarily used during the appetitive phase. The usage of the Flat syllable demonstrated genetic dependency during body sniffing, but was not significantly related to the genetic similarity score during the entirety of the mating process. In contrast, the Short syllable displayed B6-like usage consistently throughout the total duration, but not during body sniffing. This pattern corroborates previous reports of B6-like usage of the Short syllable and 129-like usage of the Chevron syllable during the total duration.^18^ Furthermore, the Two-steps syllable usage was influenced by genetic background during the appetitive state, but was not predominant. Instead, the simpler Step-up, Step-down, Down-FM, and Short syllables predominated. This aligns with earlier reports indicating that simple and short syllables (Chevron, Reverse chevron, Down-frequency modulation, Up-frequency modulation, Flat, Short) predominate during appetitive states.^13, 14, 24, 25^

Historically, it has been suggested that innate behaviors consist of two different phases: appetitive and consummatory behaviors, by ethologists^25-29^. The diverse appetitive and fixed consummatory behaviors in sexual and feeding behavior has been observed in various species, such as rodents, Japanese quail, pigeons, fish, and invertebrates^26-32^. However, there were difficulties in investigating the genetic control of two different behavioral states in those non-model animals. Thus, we used two inbred strains of mouse and their F2 progeny to figure out the genetic basis of two states. C57BL/6J (B6) and 129S4/SvJae (129) strain share same ancestry, being derived from the same subspecies *M. m. domesticus*. However, a phylogenetic analysis has shown that they have significant genetic distance^33^. Our results suggest that during the selection history of these two strains, selection and evolutionary divergence^34^ may have occurred around appetitive behaviors, whereas there was relative conservation of mounting motor pathways, possibly due to differential selection pressures such as female choice^19-20^.

Here, we contributed further insights into the genetic mechanisms underlying USV syllable patterns among inbred mice. Our findings highlight the intricate relationship between genetic background and actions in inbred male mouse courtship behavior related to USV syllable usage, emphasizing that genetic background is related to diversity in appetitive USVs but homogeneity in consummatory USVs. These insights not only enhance our understanding of genetic influences on behavioral states but also raise important questions about the specific genetic mechanisms that may govern these behaviors. Future studies could seek to directly correlate genetic background with syllable usage across actions by employing genomic sequence comparison to derive genetic scores and utilizing deep learning-based 3D pose estimation for detailed and objective behavioral analyses^35^.

## STAR METHODS

### RESOURCE AVAILABILITY

#### Lead contact

Further information and requests for resources should be directed to and will be fulfilled by the lead contact, Daesoo Kim.

#### Materials availability

This study did not generate new unique reagents.

#### Data and code availability

All data reported in this paper will be shared by the lead contact upon request. All original code has been deposited at https://github.com/sy-opennew/USV.git and is publicly available as of the publication date. Any additional information required to reanalyze the data reported in this paper is available from the lead contact upon request.

### EXPERIMENTAL MODEL DETAILS

#### Animals

All procedures were conducted according to the Korean Advanced Institute of Science and Technology (KAIST) Guidelines for the Care and Use of Laboratory Animals and were approved by the Institutional Animal Care and Use Committee. Mice (*Mus musculus*) were maintained under a 12-h light/dark cycle at 23°C with ad libitum access to food and water. Male C57BL/6J mice (B6, RRID: IMSR_JAX:000664; n=31), 129S4/SvJae mice (129, RRID: MGI:2164439; n=25), and second filial generation mice (F2, n=38) of the two strains were bred at KAIST (Daejeon, Korea). Mice were habituated on the hands of experimenters for 10 minutes each day for 5 days. The mice were then isolated in transparent housing cages for at least 7 days before courtship USV recording, and exposed to a double soundproof recording chamber (20 × 20 x 20) for 30 minutes for 3 days to minimize anxiety. Recording was done on 10- to 18-week-old male mice. For each recording, an estrous B6 wildtype female at least 8 weeks of age was introduced to the test chamber. Estrous stage was confirmed by visual inspection of the vagina^37^. B6 females were used in most of the sessions because the syllable compositions of B6 and 129 males was previously shown to be independent of the female strain^18^.

### METHOD DETAILS

#### Courtship USV induction

Female introduction was used to induce courtship USVs, since this strategy induces the production of USVs by males more effectively than other stimulants^14^. On the recording day, a male mouse was placed in one side of a transparent recording chamber for 30 minutes without recording for habituation. Then, 5 minutes of background recording was obtained and a female was introduced to the test chamber. The male mouse was allowed to investigate the female for 5 minutes (Figure 1C). Emitted USVs were recorded in the double-soundproof chamber (Figure 1B). Only data from males that exhibited proper mounting behavior and had audio recordings free of any disturbing noise was used to analyze USVs (n = number of animals; F2, n=5; B6, n=5; 129, n=4). Confounders were not controlled. The experimenter was aware of the group allocation (strain) during the experimental process, outcome assessment, and data analysis.

#### USV and courtship behavior data acquisition

USVs were recorded using a 1/4-inch microphone and amplified using a preamplifier and main amplifier (Bruel and Kjaer Inc., Denmark). Behavior video was simultaneously recorded with a video camera.

### QUANTIFICATION AND STATISTICAL ANALYSIS

#### Analysis of USV structure

Each male selected for analysis had more 100 USV syllable events (the mouse that had the least syllable events had 799 events and was F2). USV syllables were labeled using VocalMat. Produced USVs were analyzed across four time periods: the total duration, body sniffing, anogenital sniffing, and the mounting action (Figure 1B). To determine the timepoints for onset and offset of each action session, time-series video data were manually scored to onset and duration of three types of behavior: body sniffing, anogenital sniffing, and mounting. Syllables that belonged to the specific action time periods were labeled using the onset and duration data with Python. The video and audio data were synchronized using clap sound.

Probability of occurrence (PO) was adopted from general syntax analysis in Chabout et al.^36^ and calculated as:

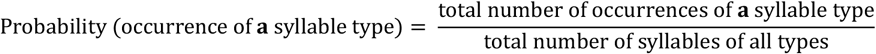

#### Generation of genetic similarity score

The genetic similarity scores of F2 individuals with respect to the B6 and 129 strains were produced using Up-FM and Chevron PO, which were reported to be majorly used by B6 and 129 mice, respectively^17,18^. The Up-FM PO x Chevron PO data points for B6 and 129 mice were fitted into a Gaussian family general linear model. The F2 data point vector was represented with the B6 and 129 regression vectors.

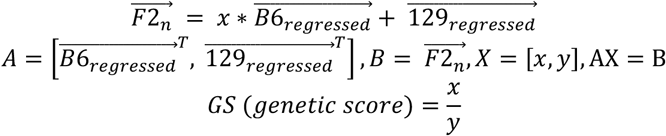

The genetic similarity score was set to be higher when the genetic background was closer to 129 mice and lower when the genetic background was closer to B6 mice.

#### Statistical analysis

The significance level was defined as p<0.05. ANOVA with two-way mixed factors was used to analyze the total duration and time period-dependent differences in vocalization data, as applied using Python 3.9.17 pingouin (0.5.3) mixed_anova. Strain was used as between-factor and syllable identity was used as within-factors (repeated measure). Tukey’s HSD test was used for pairwise multiple comparison, as applied with statsmodels (0.14.0). Sphericity test, normality test (Shapiro-Wilk), and equal variance test were done on the dependent variable (PO). All data presented in Figure 3 are given as mean ± SD.

For principal component analysis (PCA), feature matrices were created wherein each row showed a given individual mouse at each time period; each column represented one of the 9 syllables (excluding Chevron and Up-FM which were used for the genetic similarity score, and also excluding Noise), and each entry reflecting the PO of the given syllable type at the indicated time period for that individual. Data were standardized by preprocessing with StandardScaler of Scikit-learn. PCA was done using Python Scikit-learn (1.4.0) decomposition PCA for two principal components (PCs). The PC1-PC2 scatter plot was plotted in Python with color used to indicate the genetic background identity. The genetic similarity score and PC1 or PO data for each syllable were fitted into the Gaussian family generalized linear model with identity link function; constants were added to genetic similarity score before fitting.

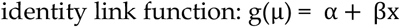

Python statsmodels (0.14.0) GLM was used to perform the fitting. Normality was tested for dependent variables (PC1 or PO for each syllable).

## KEY RESOURCES TABLE

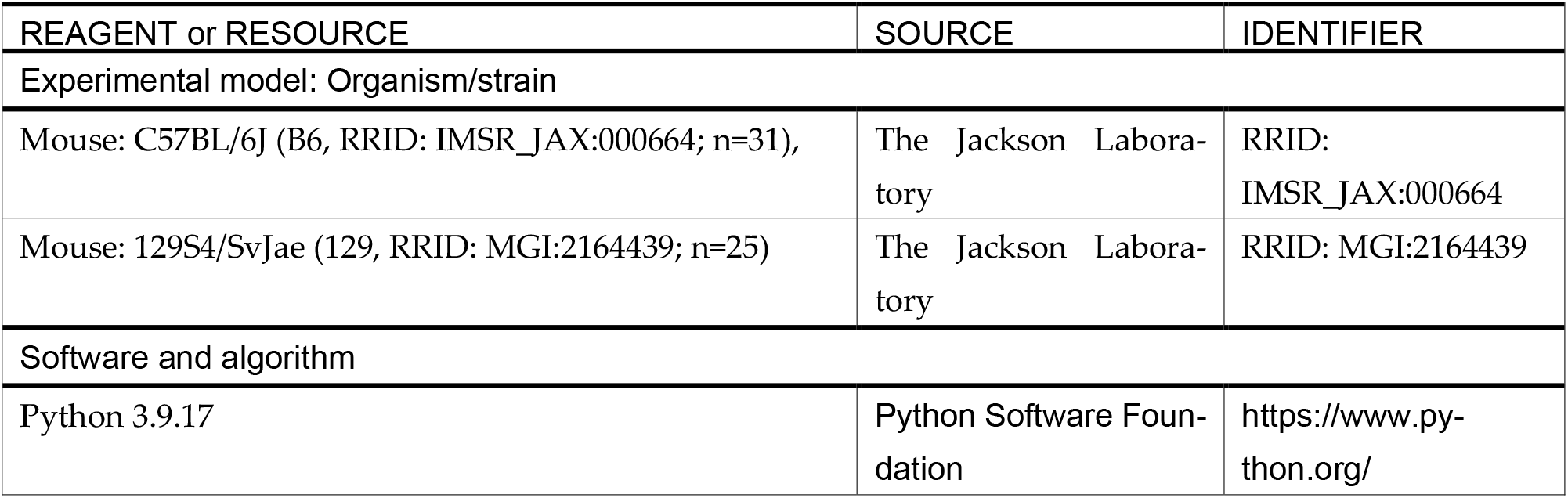

| REAGENT or RESOURCE | SOURCE | IDENTIFIER |
| --- | --- | --- |
| Experimental model: Organism/strain |  |  |
| Mouse: C57BL/6J (B6, RRID: IMSR_JAX:000664; n=31), | The Jackson Laboratory | RRID: IMSR_JAX:000664 |
| Mouse: 129S4/SvJae (129, RRID: MGI:2164439; n=25) | The Jackson Laboratory | RRID: MGI:2164439 |
| Software and algorithm |  |  |
| Python 3.9.17 | Python Software Foundation | https://www.python.org/ |

## Author Contributions

Conceptualization, J.R. and S.N.; Methodology, J.R., C.K., and S.N.; Software, S.N.; Formal Analysis, S.N.; Investigation, S.N., and J.R.; Resources, D.K.; Data Curation, S.N.; Writing—Original Draft, S.N.; Writing—Review and Editing, S.N., J.R., C.K., S.P. and D.K.; Visualization, S.N.; Supervision, D.K.; Funding Acquisition, D.K. and J.R.

## Acknowledgments

This work was supported by a National Research Foundation of Korea (NRF) grant funded by the Korean government (MSIT) (NRF-2022R1A2C3013280), the Bio&Medical Technology Development Program of the NRF funded by the Korean government (MSIT) (RS-2023-00266872), and the Basic Science Research Program through the NRF funded by the Ministry of Education (RS-2023-00249293).

## References

1. Preguica, I. M. (2017). The role of PgR+ cells in the Ventrolateral portion of the Ventromedial Hypothalamus (VMHvl) in mice female sexual receptivity. University of Coimbra. 10.13140/RG.2.2.28388.94086

2. Ishii, K. K., Touhara, K. (2019). Neural circuits regulating sexual behaviors via the olfactory system in mice. Neuroscience Research 140, 59–76. 10.1016/j.neures.2018.10.009

3. Tinbergen, N., (1951). The Study of Instinct. Clarendon Press, Oxford.

4. Georgiadis, J. R., Kringelbach, M. L., Pfaus, J. G. (2012). Sex for fun: a synthesis of human and animal neurobiology. Nat Rev Urol 9, 486–498. 10.1038/nrurol.2012.151

5. Gutierrez-Castellanos, N., Husain, B. F. A., Dias, I. C., and Lima, S. Q. (2022). Neural and behavioral plasticity across the female reproductive cycle. Trends in Endocrinology & Metabolism 33(11), 769–785. 10.1016/j.tem.2022.09.001.

6. Bialy, M., Bogacki-Rychlik, W., Przybylski, J., and Zera, T. (2019). The Sexual Motivation of Male Rats as a Tool in Animal Models of Human Health Disorders. Frontiers in behavioral neuroscience 13, 257. 10.3389/fnbeh.2019.00257

7. Moralí, G., and Beyer, C. (1992). Motor aspects of masculine sexual behavior in rats and rabbits. Adv. Stud. Behav. 21, 201– 238. 10.1016/s0065-3454(08)60145-x

8. Liu, Z. W., Jiang, N., Tao, X., Wang, X. P., Liu, X. M., and Xiao, S. Y. (2020). Assessment of Sexual Behavior of Male Mice. J. Vis. Exp. 157. 10.3791/60154.

9. Le Moëne, O., and Ågmo, A. (2019). Modeling Human Sexual Motivation in Rodents: Some Caveats. Frontiers in behavioral neuroscience 13, 187. 10.3389/fnbeh.2019.00187.

10. Ågmo, A. (2002). Copulation-contingent aversive conditioning and sexual incentive motivation in male rats: evidence for a two-stage process of sexual behavior. Physiol. Behav. 77, 425–435. 10.1016/s0031-9384(02)00874-0.

11. Chu, X., and Ågmo, A. (2016). The adrenergic a2-receptor, sexual incentive motivation and copulatory behavior in the male rat. Pharmacol. Biochem. Behav. 144, 33–44. 10.1016/j.pbb.2016.02.008.

12. Portillo, W., Antonio-Cabrera, E., Camacho, F. J., Díaz, N. F., and Paredes, R. G. (2013). Behavioral characterization of non-copulating male mice. Hormones and behavior 64(1), 70–80. 10.1016/j.yhbeh.2013.05.001

13. Matsumoto, Y. K., and Okanoya, K. (2016). Phase-Specific Vocalizations of Male Mice at the Initial Encounter during the Courtship Sequence. PLOS ONE 11(2), e0147102. 10.1371/journal.pone.0147102.

14. Wang, H., Liang, S., Burgdorf, J., Wess, J., and Yeomans, J. (2008). Ultrasonic vocalizations induced by sex and amphetamine in M2, M4, M5 muscarinic and D2 dopamine receptor knockout mice. PLoS One 3(4), e1893. 10.1371/journal.pone.0001893.

15. Heckman, J., McGuinness, B., Celikel, T., Englitz, B. (2016). Determinants of the mouse ultrasonic vocal structure and repertoire. Neurosci. Biobehav. Rev. 65, 313–325. 10.1016/j.neubiorev.2016.03.029.

16. Scattoni, M. L., Ricceri, L., and Crawley, J. N. (2011). Unusual repertoire of vocalizations in adult BTBR T+tf/J mice during three types of social encounters. Genes Brain Behav. 10(1), 44–56. 10.1111/j.1601-183X.2010.00623.x.

17. Melotti, L., Siestrup, S., Peng, M., Vitali, V., Dowling, D., von Kortzfleisch, V. T., Bračić, M., Sachser, N., Kaiser, S., Richter, S. H. (2021). Individuality, as well as genetic background, affects syntactical features of courtship songs in male mice. Animal Behaviour 180, 179–196. 10.1016/j.anbehav.2021.08.003.

18. Choi, H., Park, S. and Kim, D. (2011). Two genetic loci control syllable sequences of ultrasonic courtship vocalizations in inbred mice. BMC Neurosci. 12, 104. 10.1186/1471-2202-12-104.

19. Tam, W. Y., Cheung, K. (2020). Phenotypic characteristics of commonly used inbred mouse strains. Journal of Molecular Medicine 98, 1215–1234. 10.1007/s00109-020-01953-4

20. Atchley, W., Fitch, W. (1993). Molecular biology and evolution 10(6), 1150–1169. 10.1093/OXFORDJOURNALS.MOL-BEV.A040070

21. Fonseca, A. H., Santana, G. M., Bosque Ortiz G.M., Bampi, S., and Dietrich, M. O. (2021). Analysis of ultrasonic vocalizations from mice using computer vision and machine learning. Elife 10, e59161. 10.7554/eLife.59161.

22. Scattoni, M. L., Gandhy, S. U., Ricceri, L., Crawley, J. N. (2008). Unusual Repertoire of Vocalizations in the BTBR T+tf/J Mouse Model of Autism. PLOS ONE 3(8), e3067. 10.1371/journal.pone.0003067

23. Snedecor, G. W., Cochran, W. G. (1967). Statistical Methods (6th ed.). The Iowa State University Press, Ames, 321.

24. Portfors, C. V., and Perkel, D. J. (2014). The role of ultrasonic vocalizations in mouse communication. Current opinion in neurobiology 28, 115–120. 10.1016/j.conb.2014.07.002.

25. Hanson, J. L., and Hurley, L. M. (2012). Female Presence and Estrous State Influence Mouse Ultrasonic Courtship Vocalizations. PLOS ONE 7(7), e40782. 10.1371/journal.pone.0040782.

26. Balthazart, J., Baillien, M., Cornil, C. A., Ball, G. F. (2004). Preoptic aromatase modulates male sexual behavior: slow and fast mechanisms of action. Physiology & Behavior 83, 247–270. 10.1016/j.physbeh.2004.08.025.

27. Ball, G. F. (2008). How useful is the appetitive and consummatory distinction for our understanding of the neuroendocrine control of sexual behavior? Hormones & Behavior 53, 307–311. 10.1016/j.yhbeh.2007.09.023

28. Craig, W. (1918). Appetites and aversions as constituents of instincts. Biological Bulletin 34(2), 91–107. 10.2307/1536346

29. Sachs, B. D. (2008). The appetitive-consummatory distinction: Is this 100-year-old baby worth saving? Reply to Ball and Balthazart. Hormones & Behavior 53, 315–318. 10.1016/j.yhbeh.2007.11.015

30. Burghardt, G. M. (2020). Insights found in century-old writings on animal behavior and some cautions for today. (2020). Animal Behaviour 164, 241–249. 10.1016/j.anbehav.2020.02.010

31. Keenleyside, M. H. A. (1955). Some aspects of the schooling behavior of fish. Behaviour 8(2/3), 183–248. http://www.jstor.org/stable/4532829.

32. Paez-Rondon, O., Aldana, E., Dickens, J., Otalora-Luna, F. (2018). Ethological description of a fixed action pattern in a kissing bug (Triatominae): vision, gustation, proboscis extension and drinking of water and guava. Journal of Ethology 36, 107–116. 10.1007/s10164-018-0547-y

33. Morgan, A. P., Fu, C-P., Kao, C-Y., Welsh, C. E., Didion, J. P., Yadgary, L., Hyacinth, L., Ferris, M. T., Bell, T. A., Miller, D. R., Giusti-Rodriguez, P., Nonneman, R. J., Cook, K. D., Whitmire, J. K., Gralinski, L. E., Keller, M., Attie, A. D., Churchill, G. A., Petkov, P., Sullivan, P. F., Brennan, J. R., McMillan, L., Pardo-Manuel de Villena, F. (2016). The Mouse Universal Genotyping Array: From Substrains to Subspecies. G3 Genes|Genomes|Genetics 6(2), 263–279. 10.1534/g3.115.022087

34. Gulick, J. T. (1888). Divergent Evolution through Cumulative Segregation. Zoological Journal of the Linnean Society 20(120), 189–274. 10.1111/j.1096-3642.1888.tb01445.x

35. Kim, D., Shin, A., Jeong, Y. C., Park, S., and Kim, D. (2022). AVATAR: AI Vision Analysis for Three-dimensional Action in Real-time. Preprint at bioRxiv, 10.1101/2021.12.31.474634.

36. Chabout, J., Sarkar, A., Dunson, D. B., and Jarvis, E. D. (2015). Male mice song syntax depends on social contexts and influences female preferences. Front. Behav. Neurosci. 9, 76. 10.3389/fnbeh.2015.00076

37. Champlin, A. K., Dorr, D. L., and Gates, A. H. (1973). Determining the stage of the estrous cycle in the mouse by the appearance of the vagina. Biol Reprod. 8(4), 491–494. 10.1093/biolreprod/8.4.491.

