## Supplemental Tables for "Genetic basis of state-dependent courtship sounds in inbred mice"

Table S1. Statistics of two-way mixed ANOVA of syllables PO during total duration, body sniffing, anogential sniffing, and mounting (Strain x Syllable). *p<0.05.

| Compared group | Test used | P value | DF1 | DF2 | F | Period |
|---|---|---|---|---|---|---|
| B6 vs 129 | Two-way mixed ANOVA | <0.00002* | 8 | 56 | 6.1755 | Total duration |
| B6 vs 129 | Two-way mixed ANOVA Bonferroni correction | <0.00005* | 8 | 56 | 5.953680 | Body sniffing |
| B6 vs 129 | Two-way mixed ANOVA Bonferroni correction | <0.000006* | 8 | 56 | 7.139715 | Anogenital sniffing |
| B6 vs 129 | Two-way mixed ANOVA Bonferroni correction | 0.4076 | 8 | 56 | 1.634259 | mounting |

Table S2. Statistics of post-hoc Tukey's HSD test of syllables (Reverse-chevron; Down-FM; Flat; Short; Complex; Step-up, Step-down, Two-steps, Multi-steps) PO during total duration, body sniffing, anogenital sniffing, and mounting (Strain x Syllable).

| Figure | Comparison groups | period | syllable | Mean differences | 95% Confidence intervals | Significance level | P-values | Reject (<0.05) |
|---|---|---|---|---|---|---|---|---|
| 3A | B6 vs 129 | Total duration | Rev-chevron | -0.0006 | (-0.0057, 0.0045) | 0.05 | 0.7858 | False |
| 3A | B6 vs 129 | Total duration | Down-FM | 0.0956 | (0.0166, 0.1747) | 0.05 | 0.0243 | True |
| 3A | B6 vs 129 | Total duration | Flat | 0.0159 | (0.0043, 0.0275) | 0.05 | 0.0141 | True |
| 3A | B6 vs 129 | Total duration | Short | 0.0854 | (0.0428, 0.1281 | 0.05 | 0.0021 | True |
| 3A | B6 vs 129 | Total duration | Complex | -0.038 | (-0.0962, 0.0201) | 0.05 | 0.1657 | False |
| 3A | B6 vs 129 | Total duration | Step-up | -0.0353 | (-0.1131, 0.0424) | 0.05 | 0.318 | False |
| 3A | B6 vs 129 | Total duration | Step-down | -0.0198 | (-0.0864, 0.0469) | 0.05 | 0.5056 | False |
| 3A | B6 vs 129 | Total duration | Two-steps | -0.074 | (-0.1441, -0.0039) | 0.05 | 0.0412 | True |
| 3A | B6 vs 129 | Total duration | Multi-steps | -0.0037 | (-0.0131, 0.0057) | 0.05 | 0.3839 | False |
| 3B | B6 vs 129 | Body sniffing | Rev-chevron | -0.0008 | (-0.0068, 0.0053) | 0.05 | 0.7761 | False |
| 3B | B6 vs 129 | Body sniffing | Down-FM | 0.1019 | (0.015, 0.1888) | 0.05 | 0.0276 | True |
| 3B | B6 vs 129 | Body sniffing | Flat | 0.0116 | (0.0008, 0.0223) | 0.05 | 0.0386 | True |
| 3B | B6 vs 129 | Body sniffing | Short | 0.1076 | (0.0581, 0.1572) | 0.05 | 0.0013 | True |
| 3B | B6 vs 129 | Body sniffing | Complex | -0.0484 | (-0.1245, 0.0277) | 0.05 | 0.1764 | False |
| 3B | B6 vs 129 | Body sniffing | Step-up | -0.0515 | (-0.1322, 0.0292) | 0.05 | 0.175 | False |
| 3B | B6 vs 129 | Body sniffing | Step-down | -0.0412 | (-0.154, 0.0715) | 0.05 | 0.4157 | False |
| 3B | B6 vs 129 | Body sniffing | Two-steps | -0.0873 | (-0.1614, -0.0132) | 0.05 | 0.0271 | True |
| 3B | B6 vs 129 | Body sniffing | Multi-steps | -0.0057 | (-0.0173, 0.0059) | 0.05 | 0.2847 | False |
| 3C | B6 vs 129 | Anogenital sniffing | Rev-chevron | -0.0047 | (-0.0112, 0.0018) | 0.05 | 0.1327 | False |
| 3C | B6 vs 129 | Anogenital sniffing | Down-FM | 0.0993 | (0.0167, 0.182) | 0.05 | 0.025 | True |
| 3C | B6 vs 129 | Anogenital sniffing | Flat | 0.0223 | (-0.008, 0.0526) | 0.05 | 0.1247 | False |
| 3C | B6 vs 129 | Anogenital sniffing | Short | 0.1237 | (0.0635, 0.1839) | 0.05 | 0.0018 | True |
| 3C | B6 vs 129 | Anogenital sniffing | Complex | -0.0199 | (-0.0583, 0.0186) | 0.05 | 0.2614 | False |
| 3C | B6 vs 129 | Anogenital sniffing | Step-up | -0.0271 | (-0.0753, 0.0212) | 0.05 | 0.2262 | False |
| 3C | B6 vs 129 | Anogenital sniffing | Step-down | -0.0207 | (-0.1208, 0.0794) | 0.05 | 0.6393 | False |
| 3C | B6 vs 129 | Anogenital sniffing | Two-steps | -0.0938 | (-0.1768, -0.0108) | 0.05 | 0.0319 | True |
| 3C | B6 vs 129 | Anogenital sniffing | Multi-steps | -0.0024 | (-0.0108, 0.006) | 0.05 | 0.5254 | False |

Table S3. Statistics of generalized linear model (GLM; Gaussian family, identity link function) fitting of PC1 of 9 syllables' PO and syllables (Reverse-chevron; Down-FM; Flat; Short; Complex; Step-up, Step-down, Two-steps, Multi-steps) PO during total duration, body sniffing, anogenital sniffing, and mounting (Strain x Syllable). *p<0.05. Fig: Figure, Indep. Var.: independent variable, Dep. Var.: dependent variable, $R^2$: Pseudo R-square (CS), Bonf: Bonferroni correction, GS: genetic score.

| Fig | Indep. Var. (PO) | Indep. Var.: period | Dep. Var. | Test used | P value (α) | P value (β) | α | β | N | $R^2$ |
|---|---|---|---|---|---|---|---|---|---|---|
| 2B | PC1 of 9 syllables' | Total duration | GS | GLM | 0.0017 | 0.0005* | 2.9254 | 4.6110 | 14 | 0.5896 |

| | | | | | | | | | | |
|---|---|---|---|---|---|---|---|---|---|---|
| 2C | PC1 of 9 syllables' | Body sniffing | GS | GLM, Bonf | <0.000009 | <0.0000006* | 3.3114 | 5.2195 | 14 | 0.8592 |
| 2D | PC1 of 9 syllables' | Anogenital sniffing | GS | GLM, Bonf | 0.0025 | 0.0006* | 2.8421 | 4.4797 | 14 | 0.6329 |
| 2E | PC1 of 9 syllables' | Mounting | GS | GLM, Bonf | 0.4335 | 0.3102 | -1.6083 | -2.5350 | 14 | 0.1819 |
| - | Rev-chev | Total duration | GS | GLM, Bonf | 0.5652 | 1 | 0.007427 | -0.001627 | 14 | 0.01710 |
| - | Down-FM | Total duration | GS | GLM, Bonf | 1 | 0.9567 | 0.023204 | -0.084173 | 14 | 0.1793 |
| 4 | Flat | Total duration | GS | GLM, Bonf | 0.0561 | 1 | 0.023989 | -0.016722 | 14 | 0.1319 |
| 4 | Short | Total duration | GS | GLM, Bonf | 1 | 0.0079 | 0.015029 | -0.093976 | 14 | 0.5517 |
| 4 | Complex | Total duration | GS | GLM, Bonf | <0.00000005 | 0.0019 | 0.083358 | 0.074756 | 14 | 0.6299 |
| 4 | Step-up | Total duration | GS | GLM, Bonf | <0.00000006 | 0.5074 | 0.185463 | 0.086180 | 14 | 0.2377 |
| - | Step-down | Total duration | GS | GLM, Bonf | 0.0402 | 1 | 0.084796 | -0.001551 | 14 | 0.01133 |
| 4 | Two-steps | Total duration | GS | GLM, Bonf | <0.000000000003 | <0.00001 | 0.125039 | 0.117875 | 14 | 0.8200 |
| - | Multi-steps | Total duration | GS | GLM, Bonf | 0.0078 | 0.4501 | 0.008660 | 0.007198 | 14 | 0.2485 |
| - | Rev-chev | Body sniffing | GS | GLM, Bonf | 1 | 1 | 0.007697 | 0.000702 | 14 | 0.01204 |
| - | Down-FM | Body sniffing | GS | GLM, Bonf | 1 | 1 | 0.016806 | -0.099322 | 14 | 0.2042 |
| 4 | Flat | Body sniffing | GS | GLM, Bonf | 0.0027 | 0.0421 | 0.01876 | -0.02154 | 14 | 0.5162 |
| 4 | Short | Body sniffing | GS | GLM, Bonf | 1 | 0.1108 | 0.014752 | -0.106222 | 14 | 0.4510 |
| 4 | Complex | Body sniffing | GS | GLM, Bonf | <0.000002 | 0.0024 | 0.097691 | 0.100225 | 14 | 0.6704 |
| 4 | Step-up | Body sniffing | GS | GLM, Bonf | <0.0000000000000002 | 0.0015 | 0.208182 | 0.137533 | 14 | 0.6913 |
| - | Step-down | Body sniffing | GS | GLM, Bonf | 0.8957 | 1 | 0.084997 | -0.003098 | 14 | 0.01144 |
| 4 | Two-steps | Body sniffing | GS | GLM, Bonf | <0.00000005 | 0.0015 | 0.130363 | 0.123172 | 14 | 0.6894 |
| - | Multi-steps | Body sniffing | GS | GLM, Bonf | 0.4071 | 1 | 0.008676 | 0.007379 | 14 | 0.1515 |
| - | Rev-chev | Anogenital sniffing | GS | GLM, Bonf | 1 | 1 | 0.008513 | 0.002072 | 14 | 0.01820 |
| - | Down-FM | Anogenital sniffing | GS | GLM, Bonf | 1 | 1 | 0.027781 | 0.082423 | 14 | 0.1318 |
| 4 | Flat | Anogenital sniffing | GS | GLM, Bonf | 0.3722 | 1 | 0.045496 | -0.008531 | 14 | 0.01875 |
| 4 | Short | Anogenital sniffing | GS | GLM, Bonf | 1 | 0.0579 | 0.021029 | 0.140536 | 14 | 0.4955 |
| 4 | Complex | Anogenital sniffing | GS | GLM, Bonf | <0.000000005 | 0.0019 | 0.056422 | 0.049645 | 14 | 0.6795 |
| 4 | Step-up | Anogenital sniffing | GS | GLM, Bonf | 0.0016 | 1 | 0.114174 | 0.042965 | 14 | 0.08890 |
| - | Step-down | Anogenital sniffing | GS | GLM, Bonf | 0.3279 | 1 | 0.109010 | 0.009029 | 14 | 0.01276 |
| 4 | Two-steps | Anogenital sniffing | GS | GLM, Bonf | <0.00000000002 | <0.000002 | 0.147723 | 0.155416 | 14 | 0.8762 |
| - | Multi-steps | Anogenital sniffing | GS | GLM, Bonf | 0.0060 | 0.2416 | 0.007946 | 0.007940 | 14 | 0.3931 |
